# Achieve Handle Level Random Access in Encrypted DNA Archival Storage System via Frequency Dictionary Mapping Coding

**DOI:** 10.1101/2024.08.15.608111

**Authors:** Ben Cao, Xue Li, Bin Wang, Tiantian He, Yanfen Zheng, Xiaokang Zhang, Qiang Zhang

## Abstract

DNA as a storage medium has the characteristics of high storage density and durability, but the existing DNA storage system has a high latency, and lacks the consideration of data security. In order to improve the availability of DNA storage, this paper proposes that Frequency Dictionary Mapping Coding (FDMC) implements handle-level random access in DNA Archival storage, and a hybrid e-molecular encryption strategy and multi-level error correction algorithm are provided to ensure data security and integrity. The results of the simulation and wet experiments demonstrate that FDMC can achieve handle-level random access in lossless encrypted DNA storage systems, which balances security and convenience. In terms of read and write consistency, FDMC has significant advantages in storage density and robustness of data recovery. Even in the extreme case of DNA sequence loss of 10%, it can still recover 91.74% of the original data while ensuring storage density above 1.80 bits/nt. In summary, FDMC improves the application range of DNA as a storage medium and bridges the gap between DNA storage and traditional storage modes in the storage and reading of large-scale archives.

## Introduction

DNA is the carrier of genetic information in organisms, storing huge and complex information about life. Compared with traditional electromagnetic storage media[1, 2], DNA as a storage medium has the characteristics of high density, high durability, low energy consumption, etc. [3, 4]. With the rapid development of synthesis and sequencing technology in recent years, the use of DNA molecules [5–7] to store abiotic information has become the most concerned strategy among many organic molecular storage media[8, 9]. In 2017, Erlich et al. [10] applied fountain code in the DNA storage system and imposed constraints on single base repetition and GC content [11]. Later, Jeong et al. [12] improves the fountain code and designed the decoding algorithm of clustering and anomaly sequence clustering based on Hamming distance, which improved the base utilization rate of 6.5-8.9%. Anavy et al. [13] adds logical redundancy to the fountain code technology and used complex bases to encode 2.12MB files, which increased the coding density by 24% compared with Elich’s work. In addition to the fountain code, WMU code [14], HEDGES[15] and RS code [8, 16] have also made certain achievements in DNA storage. Ping et al. [17] proposes Yin-yang codec in 2022, which realized the approximate physical storage density of theoretical values, but the above method can only perform sequential accesses during data access.

The high latency during the read-write process imposes limitations on the rapid implementation of DNA storage. This latency is evident not only in the synthesis and sequencing processes but also in the data read-write operations. Random access allows direct access to any location within the storage medium as needed, effectively reducing the latency issues associated with sequential access. Currently, achieving random access in DNA storage primarily relies on PCR[18–20]. Yazdi et al. [20] write 17KB data into DNA and selectively edited three pieces of information, which is the first to realize random storage and selective rewriting. However, the length of the DNA sequence used in the stored process exceeded 1000bp, and long DNA sequences are more difficult to synthesize relative to short sequences around 200bp. Addressing schemes [21] enable random access to stored data in large-scale DNA storage systems. However, the random access achieved is between files and cannot specifically extract particular information within a single file. Due to the limited efficiency of single-primer PCR and the constraints of large libraries of carefully designed orthogonal PCR primers on DNA storage capacity and scalability, the nested primer address system [19] is proposed. However, it requires more complex experimental protocols and costlier reagents to achieve high specificity, as well as a greater DNA address space. Therefore, Winstond et al. [22] report a combinatorial PCR method that exhibits excellent specificity in retrieval. Thermal-sealed polymerase chain reaction [23] enables multiplexing and repetitive random-access partitioning of DNA files based on thermally meltable semi-permeable membrane microcapsules. Compared to repetitive random access, the platform’s performance exceeds that of non-partitioned DNA storage and reduces amplification bias by tenfold during multiple PCR processes. In terms of PCR-free random access, a DNA microdisk [24] for DNA data management is proposed, which exhibits excellent indexing capabilities. Utilizing other inorganic materials for random access is also a feasible solution, whereby encoding data’s DNA file sequences are encapsulated within silicon dioxide spheres labeled with single-stranded DNA barcodes [25], enabling random access to files by selecting barcodes. However, the above storage schemes that support non-sequential reading still have problems with low utilization of storage bases and low retrieval efficiency[26] [27].

Although DNA storage technology has advanced to the point where the entire process can be implemented in the laboratory, there are still unsolved problems with DNA storage. On the one hand, the existing DNA synthesis and sequencing technologies lead to high cost and high latency of DNA storage reading and writing. The high latency of reading data in DNA storage is not only reflected in the delay of the current sequencing technology, but also reflected in the existing data encoding and reading methods. For example, in DNA fountain code, Erlich et al. [10] use a large amount of redundancy to ensure data integrity, and the stored files also need to be fully sequenced and put into the decoder before decoding. On the other hand, the security of DNA storage lacks assurance, and although there are considerations of data security through XOR[28], DNA secret key[29], etc., these information security schemes cannot quantify data security, and there is still room for improvement in data security in the AES secret key construction scheme based on human STR proposed by Grass et al.[30]

In order to solve the above problems in DNA storage, this paper proposes a DNA archival storage system based on frequency dictionary mapping coding (FDMC) from the perspective of improving data security and reading integrity and efficiency (Figure 1a). It implements handle-level random access in large-scale text data and verifies the significant advantages of FDMC in storage density and robustness of data recovery through experiments (Table 1, Figure 2). As shown in Figure 1b, FDMC is encoded by frequency dictionary satisfying constraints, which not only guarantees no homopolymer and balanced GC content at the source, but also reduces the large amount of redundancy generated by binary as intermediate code, thereby improving coding efficiency and reducing error rate. After encoding, complete data encoded as DNA sequences are split into short DNA sequences in an overlapping way (Figure 1c), which provides backup for error-prone bases at the beginning and end, and improves the robustness of data recovery. Compared with the current advanced coding methods, FDMC can effectively ensure the absence of no homopolymer and balanced GC content, and meet the data security requirements to maintain storage density above 1.80 bits/nt. During data storage, the complete dictionary is encoded in the circular plasmid, and the data is encoded in the DNA pool (Figure 1a). Since file size and word frequency are not necessarily linked (Supplementary file Fig. 2), the effect of different text files on plasmid size is weak. In addition, in order to ensure the security and integrity of the stored data, FDMC also supports the hybrid e-molecular encryption strategy and multi-level error correction algorithm. Figure 1e and Figure 5, respectively, show the key construction process and error correction algorithm. The results of silicon-based and vitro experiment show that FDMC not only achieve lossless and secure storage of text data, but also can realize handle-level random access in DNA archival storage systems (Figure 1d, Figure 4), which balances security and convenience.

**Figure 1.**
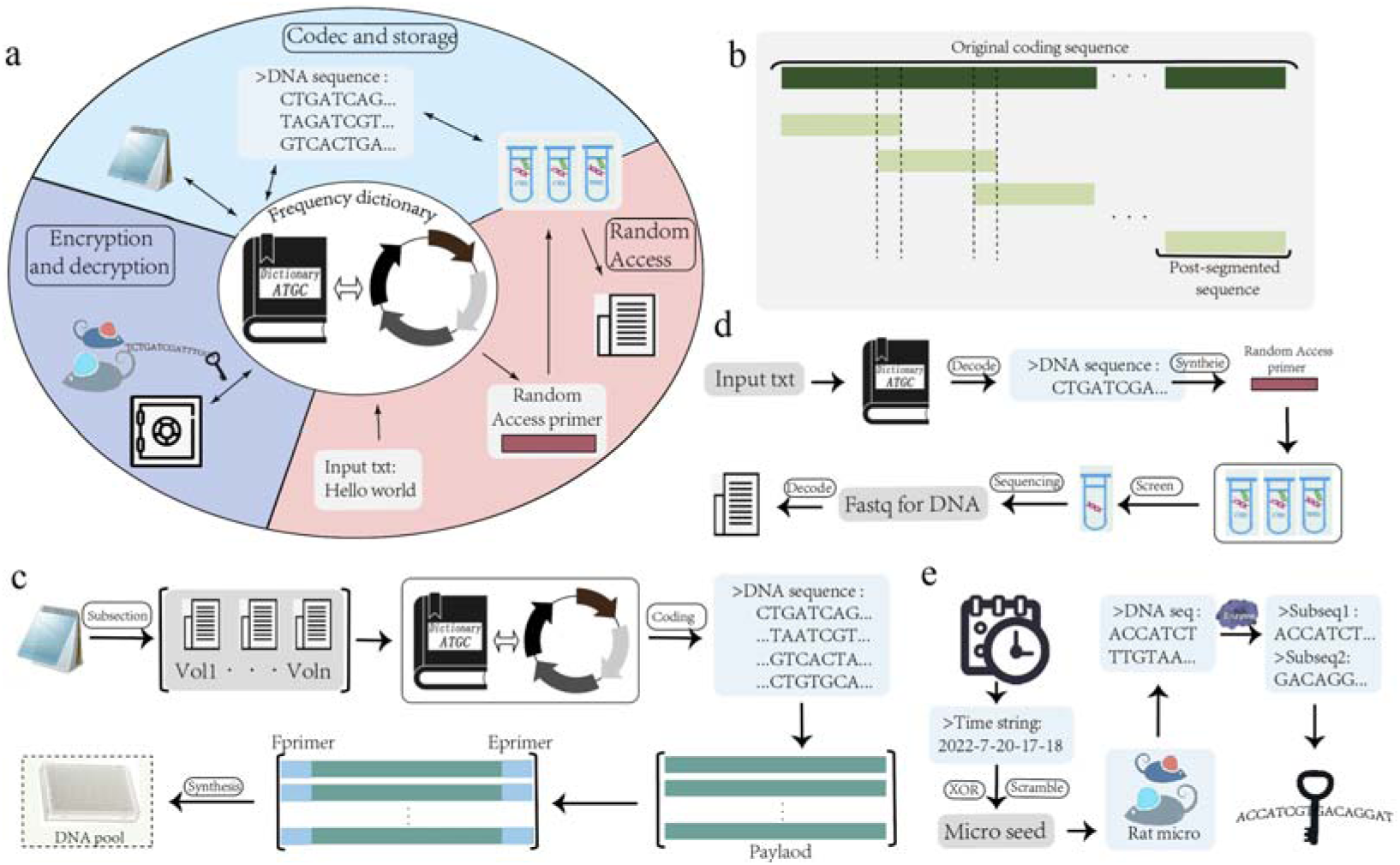
DNA archival storage system based on frequency dictionary mapping coding. (a) shows three important modules in FDMC, codec and storage, random access, encryption and decryption. (b) is the detailed process of frequency dictionary mapping coding. Through frequency dictionary satisfying constraints, text data are encoded as DNA sequences by FDMC, and then primers are added to be stored in DNA pool. The splitting of encoded DNA sequence into short DNA sequences is carried out by overlapping splitting in (c), which can add backup to the beginning and end of error-prone DNA sequences and improve the robustness of data recovery. (d) is the handle level random access process. By synthesizing random access primers, the target DNA sequence is extracted by PCR and purification, and the target data is obtained by sequencing and decoding. (e) shows the key generation process of encryption and decryption, and constructs the AES-128 encryption key with high key entropy through time series and mouse microsatellite to protect the security of data.

**Figure 2.**
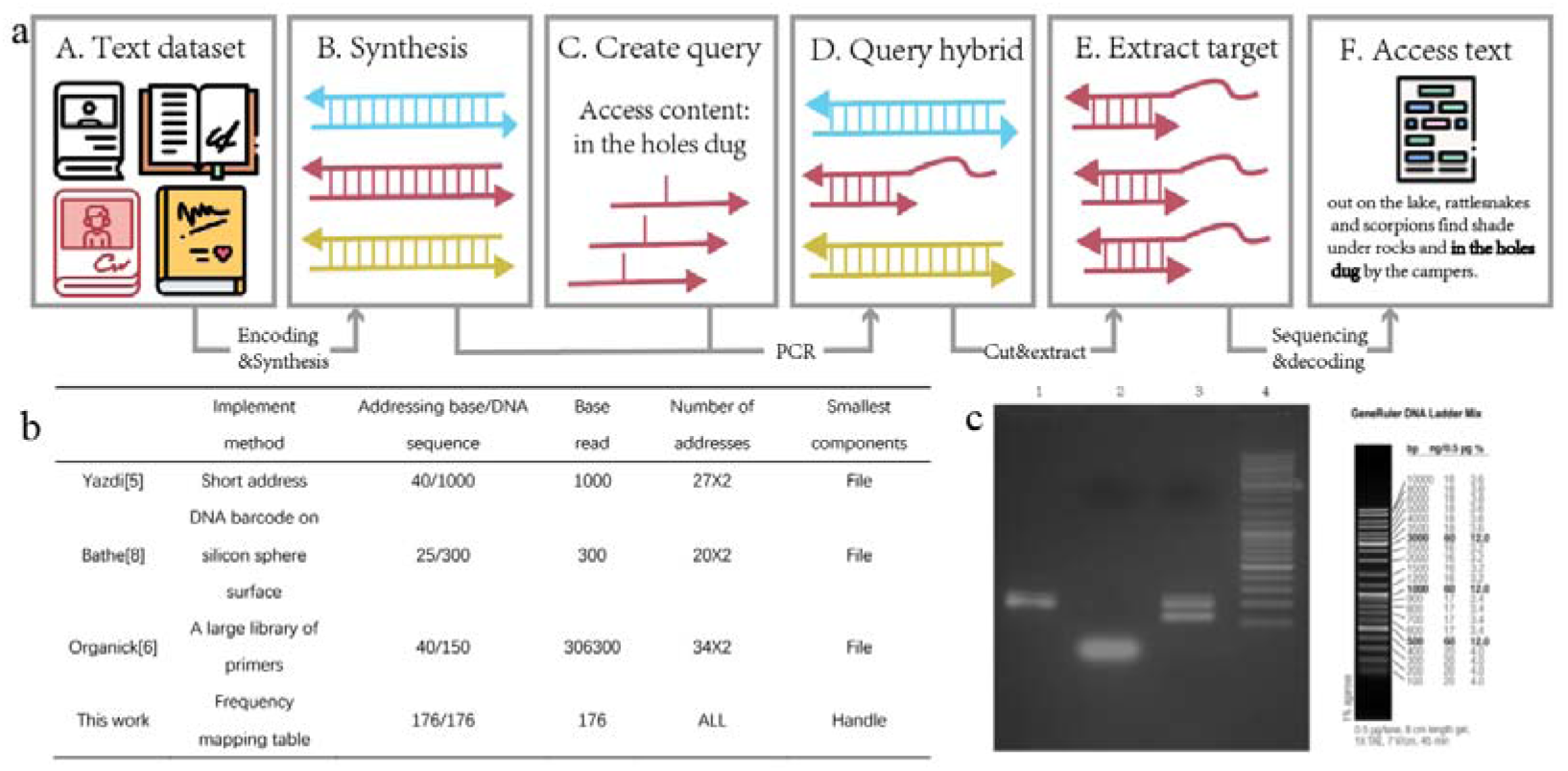
FDMC implements the whole process of handle level random access. (a) is the whole process of implementing handle-level random access in FDMC. When access is needed, random access primers are constructed to achieve hybridization between random access primers and DNA target sequences in DNA storage database, and access to target data is achieved after extraction, purification, sequencing and decoding. (b) is the comprehensive comparison of random access capability in DNA storage system. The first row of the table represents the implementation method of random access respectively. The Smallest components refer to the smallest unit that can be accessed randomly, and the Number of addresses refers to how many additional high quality oligonucleotides are required by the current storage system to act as address bits. Addressing base/DNA sequence refers to the percentage of random access functional bits to the entire oligonucleotide sequence. Base read refers to the minimum number of bases that need to be sequenced and decoded when a sentence or phrase needs to be accessed. It can be seen from the results in the table that FDMC can not only realize random access, but also can act as the addressing target with more bases, which can realize efficient random access in DNA storage at a lower cost. The results of agarose-gel electrophoresis after PCR were shown in (c). It can be clearly seen that the target product is obvious in the third pass. Compared with mark, it is verified that the length is in the expected range and the distance is far away, which is easy for gel recovery and purification.

**Figure 3.**
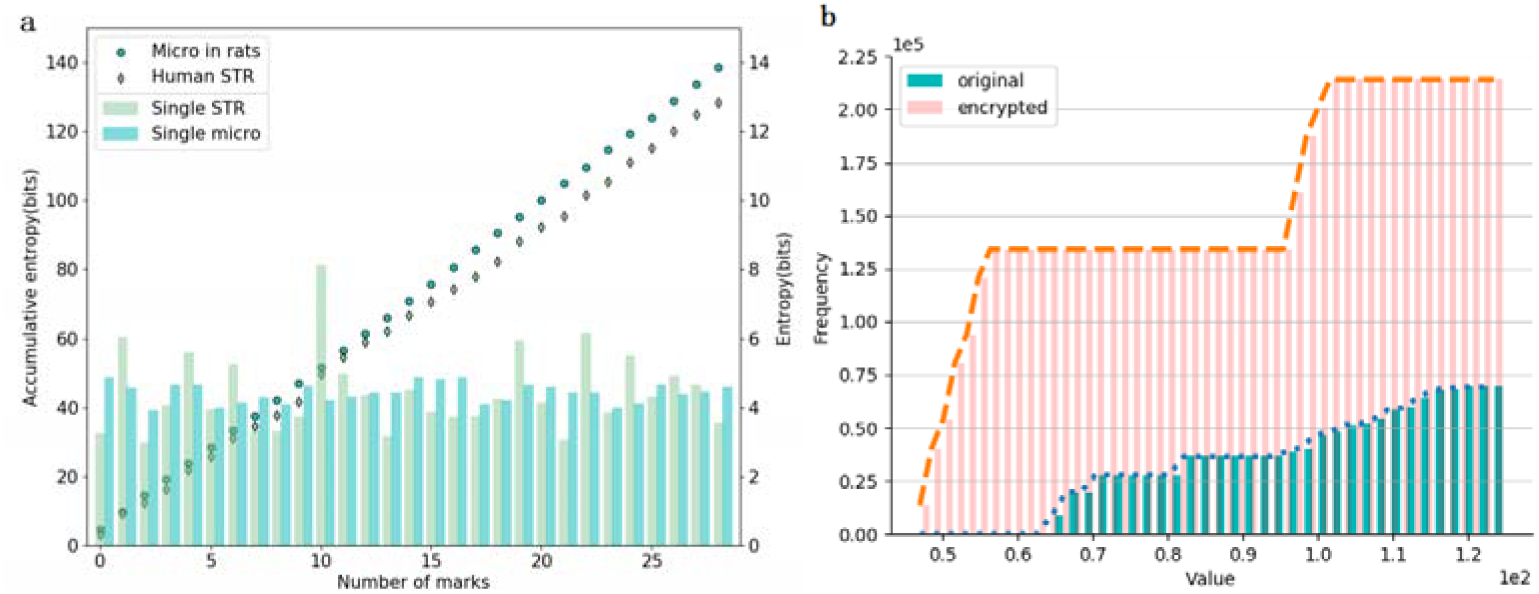
Comparison of AES-128 key performance generated by Micro of rats and Human STR. Figure 3a respectively compares the key entropy of a single Micro of rats and Human STR, and the Micro of rats is stable. Moreover, the sum key entropy of Micro of rats and the results obtained by Grass et al under the same calculation method are also compared. Figure 3b is the cumulative histogram of plaintext (blue) and ciphertext (orange) ASCII codes. Flatter ciphertext histograms can effectively hide statistics.

**Figure 4.**
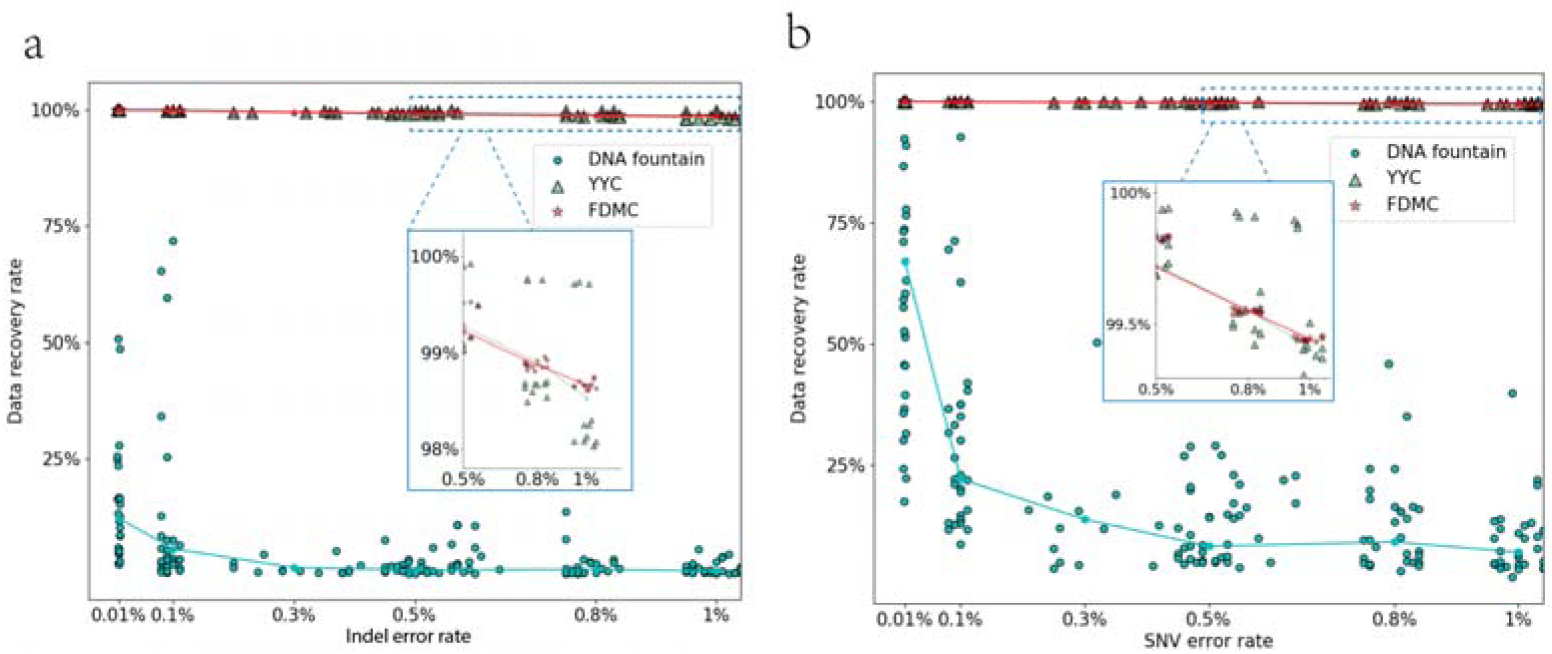
Robustness analysis of FDMC and DNA fountain and YYC under insertion error. (a) and SNV error (b). The horizontal axis of Figure 4a and 2b respectively represents the insertion error rate and SNV error rate, and the vertical axis represents the data recovery rate. Blue circle, green triangle and red five-pointed star represent DNA fountain code, YYC and FDMC respectively, and line segments of the same color are the median lines of the current error rate. The results in the figure show that YYC and FDMC have significant advantages over fountain codes. Moreover, it can be seen from the enlarged figures of Figures 2a and 2b that the variance of recovery rate of YYC is larger and more discrete, and the recovery rate of YYC is more stable. Moreover, in most cases, the line segment representing FDMC is above YYC, indicating that FDMC has higher high robustness than YYC.

**Figure 5.**
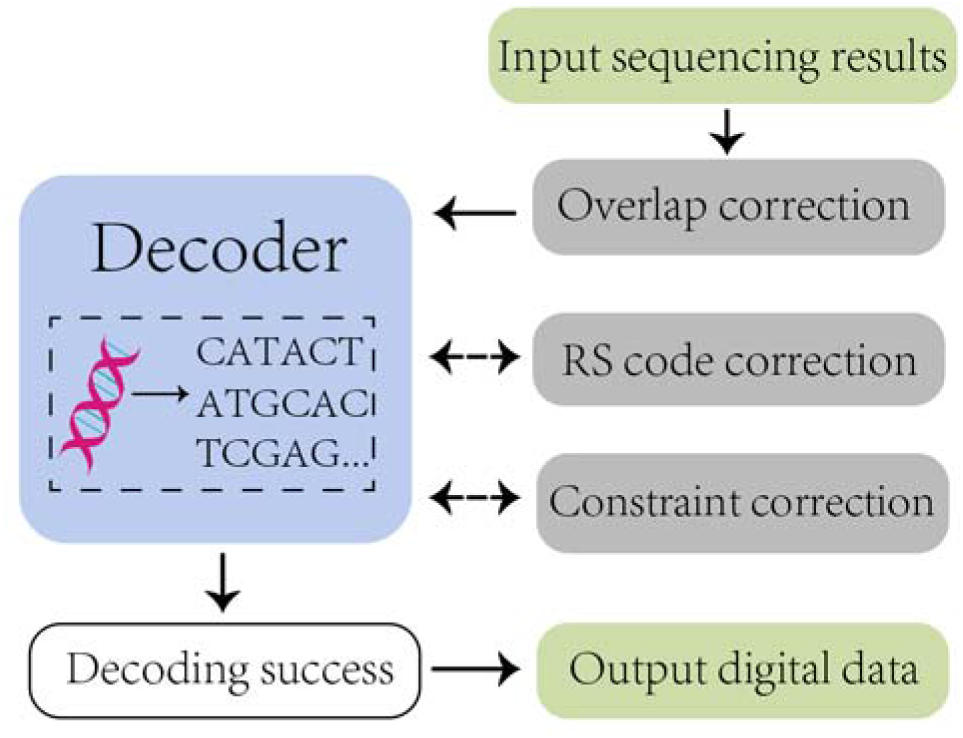
Schematic diagram of multi-level error correction scheme. The gray part shows three different levels of error correction methods. First, the sequencing error of the current part is corrected by overlapping part, and then the decoder is used to determine whether the decoding is successful. If the next level of RS code error correction is performed without fully decoding the data, the RS code of GF (28) in this paper can correct four bases on the current sequence. Continue through the decoder after completion and output digital data if successful. If there are still errors in the last level of constraint error correction, the error correction ability of the multistage error correction scheme is far greater than the theoretical DNA storage error rate (Figure 4, Table 2).

**Table 1.**
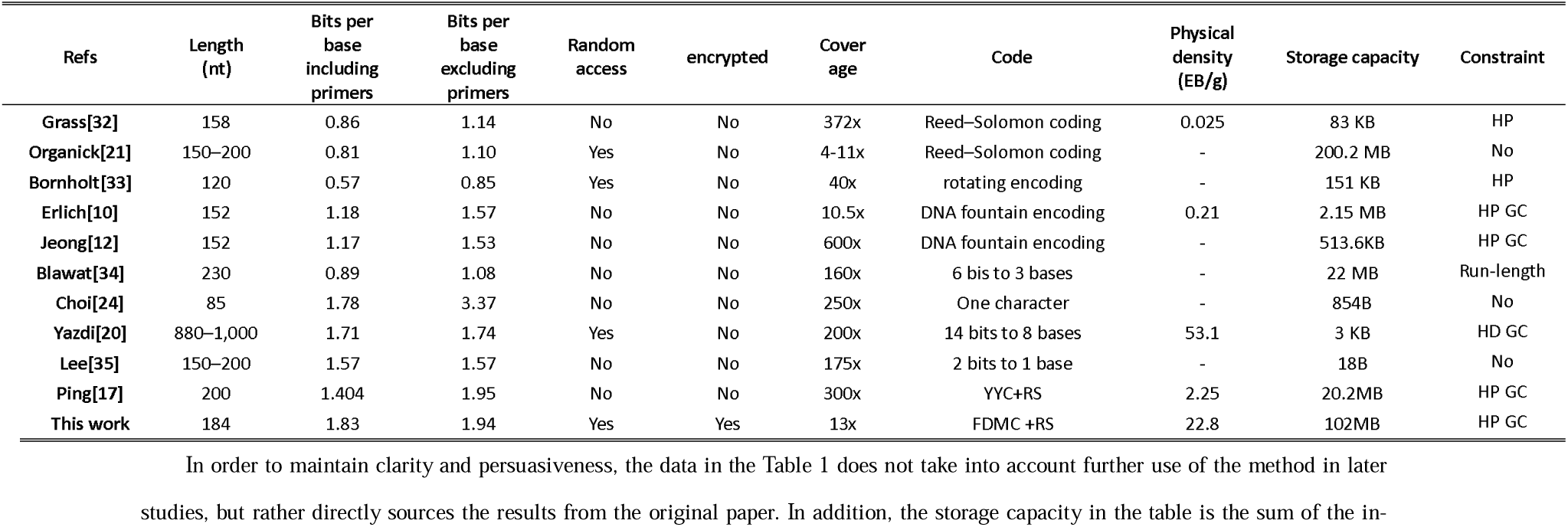
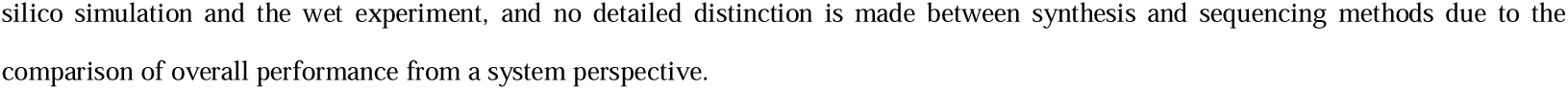
Comparison of performance indicators between FDMC and other advanced DNA storage schemes. The first line Refs represents the proposer of the scheme, the Length represents the length of the DNA sequence used in the DNA storage scheme, two cases of storage density with and without primers, namely Bits per base including primers and Bits per base excluding primers, and compares whether random access and encryption methods are supported. Coverage represents the sequencing coverage when read data, which can reflect the robustness of coding scheme. Code and Constraint represent the codec scheme and the constraint type that is satisfied, respectively. At the same time, the physical storage density and storage capacity are also compared. The calculation method is shown in the steps in supplementary file 1, and the results in the table show that FDMC has significant advantages in both storage performance and security.

**Table 2.** Comparison of sequence loss rate of FDMC with fountain code and YYC at base loss rates of 1-10%. The first row represents the deletion error rate respectively. The results in the table show that the recovery rate of YYC and FDMC is significant relative to fountain code, and FDMC still has certain advantages relative to YYC recovery rate. In each case, it is the scheme with the highest data recovery rate among the three.

|  | 0.01 | 0.015 | 0.016 | 0.017 | 0.018 | 0.019 | 0.02 | 0.03 | 0.04 | 0.05 | 0.06 | 0.07 | 0.08 | 0.09 | 0.1 |
| --- | --- | --- | --- | --- | --- | --- | --- | --- | --- | --- | --- | --- | --- | --- | --- |
| Fountain[10] | 100.0<br>% | 100.0<br>% | 100.0<br>% | 100.0<br>% | 84.66<br>% | 53.20<br>% | 9.82%<br>% | 7.84%<br>% | 4.82%<br>% | 2.90%<br>% | 2.18%<br>% | 2.04%<br>% | 1.76%<br>% | 1.68%<br>% | 1.42%<br>% |
| YYC[17] | 99.00<br>% | 98.50<br>% | 98.40<br>% | 98.30<br>% | 98.20<br>% | 98.10<br>% | 98.00<br>% | 97.00<br>% | 96.00<br>% | 95.00<br>% | 94.00<br>% | 93.00<br>% | 92.00<br>% | 91.00<br>% | 90.00<br>% |
| FDMC | 99.17<br>% | 98.76<br>% | 98.68<br>% | 98.60<br>% | 98.51<br>% | 98.43<br>% | 98.35<br>% | 97.52<br>% | 96.70<br>% | 95.87<br>% | 95.04<br>% | 94.22<br>% | 93.39<br>% | 92.57<br>% | 91.74<br>% |

## Result

### General principles and characteristics of FDMC

Storing information in DNA is the latest technology for data storage, but an old fashion existing in nature. Moreover, since it constitutes the human genome, the technology to read it will never be outdated, unlike the reading software in electromagnetic storage media [31]. Based on this characteristic, this paper proposes a long-term file storage scheme using DNA as a storage medium. The general principle of FDMC is to map words to elements in the filtered DNA frequency dictionary. Since oligonucleotides in the frequency dictionary meet the combination constraints, the encoded code word does not show homopolymer and extreme GC content. In the first step, oligonucleotides with a length less than ten were screened by GC content, no homopolymer and spacer constraint, and the oligonucleotide sets were obtained by ascending sequence of oligonucleotide lengths. Then, word frequency statistics were performed on the stored text data, which were arranged in descending order according to the occurrence frequency and combined with the oligonucleotide set to form a mapping dictionary. Finally, the data will be encoded according to the mapping relationship between words and oligonucleotides, and real-time pre-encoding verification of encoded-words will be conducted to determine whether there is ambiguity (multiple decoding results). If there is ambiguity, fine-tune oligonucleotides of the same length in the mapping dictionary until there is no ambiguity in the encoding result of the text file.

In this chapter, we first analyze FDMC from a systematic perspective, focusing on key metrics such as encoding density, random access, and encryption et.al. By comparing it with other advanced methods, we can demonstrate the comprehensiveness of FDMC as a lossless DNA storage system. Notably, FDMC is the only DNA storage system with handle-level random access capability and encryption methods. Secondly, we evaluate FDMC’s random access performance through a combination of dry and wet experiments. Figure 4 illustrates the process and results of handle-level random access, showing that FDMC achieves satisfactory results in terms of both address bit count and access precision. Additionally, encryption is one of FDMC’s key features. Building on the work of Grass et al. [30], FDMC constructs a more secure key to encrypt the frequency dictionary using rat microsatellites and time variables, ensuring the security of DNA stored data. More importantly, encrypting the dictionary does not affect the handle-level random access process. DNA storage is currently mainly used for cold data, which may experience base mutations or sequence loss in long-term storage or special environments. Therefore, it is necessary to design robustness experiments in the absence of error-correcting codes for the storage system. Besides comparing data recovery rates under a general 1% error rate, we also designed more competitive sequence loss experiments. The results show that even in the extreme case of a 10% DNA sequence loss, it can still recover 91.74% of the original data while ensuring a storage density above 1.80 bits/nt.

### Comprehensive analysis of DNA storage systems

In order to illustrate the application potential of FDMC in actual storage, this paper selects more than 100 English classics to be encoded through FDMC. By analyzing the statistical data during the coding process, it is shown that FDMC has no fixed preference in terms of data structure, and the inevitable relation between word frequency and file size is excluded (supplementary file Figure S2), so the increase in storage files does not significantly lead to dictionary size expansion, i.e., plasmid size is essentially constant. To further measure the encoding performance and storage capacity of FDMC, the relevant performance indicators were compared with other advanced DNA storage systems (Table 1). Common indicators to verify the performance of storage systems include storage density, storage capacity, etc. However, in DNA storage systems, a comprehensive evaluation should be carried out on sequencing coverage, access mode, DNA sequence length, constraints met, and physical density. Storage density is a measure of how much data can be stored per base, but current DNA storage systems can calculate storage density in a variety of ways, such as net information density, Bits per base including primers, Bits per base excluding primers, etc. However, this paper believes that storage density should be calculated by dividing data size by all bases used, i.e., Bits per base, including primers. But in the interest of fairness and respect for the work of those who came before, this paper compares both Bits per base, including primers, and Bits per base, excluding primers. The increase in coverage will make it more difficult to read information, and the storage capacity can reflect the size of data that the DNA storage system can hold. In addition, the difference in coding mode has a great impact on the performance of the DNA storage system, and the coding result can meet the constraints to reduce the occurrence of errors.

Due to the error-prone and difficult synthesis of ultra-long DNA sequences in the process of synthesis and sequencing, DNA sequences less than 200 lengths are still used as the minimum storage structure in this work. It can be seen from Bits per base that the FDMC proposed in this paper achieves a higher storage density, which is more than 20% higher than the previous optimal value and closer to the theoretical optimal value. This is because FDMC adopts frequency mapping dictionary encoding mode, which has a certain data compression ability and improves base utilization rate. In terms of stored data security, FDMC supports a highly competitive hybrid e-molecular encryption strategy, which overcomes the defect of no security module in other schemes. In FDMC, only the frequency dictionary can be encrypted to complete the encryption of all stored information, which improves the compatibility with traditional information security protection methods and thus improves the availability and security of DNA storage. In the aspect of data reading, FDMC can realize handle-level random access, compared with sequential reading, which can reduce the latency of reading data.

In addition, compared with the address bit and primer-based approach proposed by Organick [21] and Bornholt [33], the FDMC’s random access method based on frequency mapping coding has the characteristics of low time complexity and can realize the handle level random access. Because all the elements in the frequency mapping dictionary of FDMC satisfy the no homopolymer and GC content constraints, the encoding result of FDMC also satisfies the above constraints. Other constraints such as Run-length, Hamming distance constrains (HD), etc. are also discussed in the supplementary document 2. Coding results that satisfy constraints can reduce read/write error rate and thus sequencing coverage. Sequencing coverage is also an important index in evaluating DNA storage systems. The sequencing coverage of FDMC is slightly higher than that proposed by Erlich et al. [10], but it is even an order of magnitude lower than all other schemes. Lower coverage indicates not only stronger coding robustness, but also lower error rates. Physical density refers to the amount of effective information that can be stored by a storage medium per unit weight. Relative storage density is better reflected in the physical space occupied by the information stored in actual applications. The physical storage density of a DNA storage system is usually calculated as a base pair weight equal to approximately 650 Daltons, as detailed in the supplementary document 1. As can be seen from Table 1, the physical density of FDMC is only slightly lower than that of Yazdi et al, but the storage capacity of Yazdi’s scheme is only 3kb, which lacks practicability. In addition, FDMC is significantly superior to all other schemes and remains the same order of magnitude as Yazdi’s physical density [20] for high storage capacity. In terms of storage capacity, FDMC is significantly superior to other schemes and remains in the hundred MB range with Organick [21], further demonstrating FDMC’s storage performance.

### Handle level random access performance verification

The high latency of reading data in DNA storage is reflected not only in the latency of current sequencing techniques but also in the existing way data is encoded and read. Random access in computer systems refers to arbitrary access to a specific block of data, while sequential access requires traversing all data. However, random access is now common at the computer system level. However, at the hardware level, random access storage is still a relatively advanced access method. For example, in traditional HDD, because HDD is a mechanical hard disk, the location data needs seeking and rotation to locate the data block to be read and written, which cannot complete random access in a true sense [36]. However, SSD directly calculates the location of data storage by maintaining mapping tables. Therefore, SSDs have real random access capability. Similarly, FDMC proposed in this paper also completes random storage by maintaining mapping tables (frequency dictionary). In order to illustrate the unique advantages of FDMC in random access, in vitro silicon-based simulation and wet experiments were carried out in this section, as shown in Figure 4.

To determine the compatibility of FDMC with current biochemical techniques (DNA synthesis, PCR amplification, assembly techniques, DNA sequencing), as well as random access efficiency in real-world biological experiments. The text data was encoded into a DNA sequence by FDMC, synthesized into a DNA strand, and stored appropriately. Take “in the holes dug” as an example, access the DNA database, and apply the same FDMC code to the phrase to be accessed. The query ssDNA is then created and amplified by PCR after mixing ssDNA with the DNA database. After PCR, agarose gel electrophoresis was performed, and the PCR products were recovered and purified using a gel recovery kit to obtain the purified products and then sequenced and decoded to obtain the target text. In Figure 4c, the first line is the DNA database mixture, the second line is a random access primer that encodes the phrase, the third line is the reaction product, and the fourth line is the mark. The target product is obvious in the third track, and the comparison with the mark verifies that the length is in the expected range. Moreover, it can be seen from the sequencing results (supplementary material 1 & Figure S3) that the access target is consistent with the expected results, and the handle level random access in DNA archival storage is realized after sequencing and decoding.

In Figure 4b, the random access performance of FDMC is quantified by comparing and analyzing with the advanced random access DNA storage system[20, 21, 25]. In terms of the implementation of random storage, compared with other methods using address bits [20, 21] or barcodes [25], FDMC can realize handle-level random access through frequency mapping table. Although quaternary (ATGC) has more permutations and combinations than binary (0,1), the fault tolerance rate of address bits is lower than that of data bits [37], so many biological constraints need to be met to ensure the accuracy, such as GC content, no homopolymer, HD, etc. In general, when large-scale data is used for DNA storage, the set of high-quality address bits is still scarce. That’s one reason why Yazdi[20] and Bathe[25]can only store data on a small scale and why Organick [21] can only get file-level random access.

Improving the random access capability can be achieved by expanding the address bit length to meet the constraint of the address bit set, but this increases the physical redundancy and reduces the base utilization. Addressing base/DNA sequence showed that addressing base accounted for a large proportion of the whole DNA sequence in previous works, especially the addressing redundancy in Organick’s work [21] which accounted for 26.7% of a single DNA sequence, which was not conducive to giving full play to the potential of DNA molecules as storage media. All data bits in FDMC can be used as addresses, which realize the data and address function of the same oligonucleotide simultaneously. Base read refers to the minimum number of bases that need to be read and decoded to access any random sentence in a DNA storage system. For example, in Organick’s large-scale storage system [21], accessing a sentence in a Text requires 306300bp base reading and decoding, which undoubtedly increases the time complexity and decoding cost. However, in FDMC, because of the frequency mapping table, each encoding block can be addressed through the mapping table so that a certain sentence or phrase can be accurately accessed. Compared with Bathe et al. [25], who had the least bases read, FMDC can still reduce the minimum number of bases in a single access by nearly double, greatly reducing the read and write latency.

### Encryption security evaluation

The encryption strength mainly depends on the encryption mode and the complexity of the key, and the length of the key is the decisive factor of the complexity of the key. Biometric key generation is an emerging key design method, and STR maps are relatively mature in paternity testing due to their high recognition, and researchers have prelim natively demonstrated the potential of using human STR maps to generate AES keys [30]. In the paternity test, about 20 STR loci (autosomal 22+ gender chromosome) are commonly detected [38]. Since the common human genome is diploid, the detection of these STR loci is usually sufficient to determine the relationship, but this may cause trouble when the STR map acts as the key. As shown in Figure 3a, the round dots and the light green bars represent Grass [30]. When only 17 STR sites can be used to obtain the key with an entropy of 80, 20 years have passed since the cracking of the 64-bit key. It is recognized that the key entropy must be greater than 100 before it is recognized that the supercomputer will not crack it by force, and it is dangerous to use the key with a key entropy less than 100. From the perspective of considering side-channel attacks [39], using only human STR increases the possibility of obtaining prior knowledge.

In order to improve the key complexity and reduce the risk of attack, this paper uses more microsatellites of diploid organisms as the key generation seed of the AES-128 encryption algorithm and adds time parameters and biochemical reaction operations to improve the difficulty of brute force cracking and side-channel attacks. It is clear from the points in Figure 3a that the sum of entropy in the Micro of rats (calculated in the same way as Grass[30]) is consistently better than that in the human STR map alone, and the effect becomes significant after the number is greater than 10. In addition, considering the entropy of individual seeds, although some Human STR is better than Micro of rats, the entropy of Micro of rats is greater than that of Human STR in most cases. According to the barrel theory, the security of the encryption system depends on the shortest piece of wood. The mean-variance of Human STR is larger than that of Micro (supplementary Table S1), indicating that selecting different Human STR as key seeds will lead to large fluctuations in key entropy, which is not conducive to encryption performance. This is because the key entropy, in turn, determines the amount of guesswork required for brute force cracking because the encryption system often needs to consider the number of attempts in the worst case, so the key entropy with large variance will have an impact on the security of the entire encryption system. As can be seen from the bar chart in Figure 3a, compared with the key entropy of Human STR, the Micro of rats not only adds up to a larger sum when the number of digits is the same but also has a smaller variance of the key entropy, indicating that different Micro of rats as key seeds have no significant influence on the key entropy.

To better illustrate the data security performance of FDMC, this section also uses a histogram (Figure 3b) to measure the hiding effect of ciphertext on statistics. However, when features in the text cannot get all possible values, many 0 values will appear in the common histogram. These 0 values will affect the intersection operation of histograms so that the matching values cannot correctly reflect the distribution differences between different texts. Therefore, cumulative histograms are used in this paper to reflect the distribution differences of texts more clearly. Figure 3b shows the cumulative histograms of plaintext and ciphertext ASCII codes. The plaintext histograms are more volatile, while the ciphertext histograms are flatter, which hides the statistics in plaintext well and reduces the possibility of DNA stored data being attacked by side channels.

### Robustness analysis of data recovery

High precision read and write consistency is a necessary condition for DNA storage systems, but both the process of data writing and the process of base decoding to data will introduce some errors. These errors fall into two main categories. One is due to the inherent errors introduced by synthesis and sequencing technology and equipment, such as the synthesis error of a small number of bases, which can be corrected by increasing the sequencing depth. The other is errors that occur in all DNA sequences, such as insertion, deletion (indel), and substitutions (single-nucleotide variations, SNV) errors during synthesis and PCR, but for which increasing sequencing coverage is of little significance. In particular, insertion and deletion errors can change the length of DNA sequences, and error diffusion caused by these error types can be fatal to the data stored by DNA [27]. What’s more, in extreme cases, the entire DNA sequence of stored information will be lost. These systematic errors [17] bring challenges to stored procedures. So, in this section, the data recovery robustness of FDMC, DNA fountain and YYC was tested for base error without error correction code and sequence loss with error correction method.

### Base error without error correcting code

Although FDMC is a lossless DNA storage coding system with multi-layer error correction function, in order to verify the robustness of FDMC in the face of complex errors, this paper analyzes the data recovery rate without introducing external error correction (RS, etc.). The comprehensive error rate are introduced into the DNA sequence in a gradient within the range of 0-1% same as Heckel et al. [40]. The data recovery rates of two error types, indel errors, and SNV errors, were respectively compared with the most representative scheme (DNA fountain) [10] and the latest scheme (YYC) [17].

The results show that compared with fountain code and YYC, FDMC can better deal with indel and SNV errors, and the data recovery rate can keep above 98.5% and 99%, respectively. In the Figure 4, the results of FDMC and YYC [17] are very similar because when the error rate is less than 1%, both FDMC and YYC can better complete the original data recovery, and the recovery rate is close to 100%. However, it can be seen from the enlarged details of Figure 4 that the distribution of FDMC scatter points is more uniform, indicating that FDMC data recovery capability is more stable. Moreover, the first function of FDMC fitting scatter is higher than YYC, indicating that FDMC has a stronger data recovery ability in the face of indel and SNV errors. As can be seen from Figure 4, compared with YYC and FDMC, DNA fountain has a significant disadvantage in data recovery because the DNA fountain encoding algorithm requires many XOR, which will further improve the correlation between data blocks. So, when a system error occurs, it is irretrievably propagated to other data blocks. Although the RS error-correcting codes were introduced in Erlich’s work [10] by adding redundancy, the RS and other error-correcting codes have certain limitations in error-handling ability [41]. When the error handling ability is exceeded, some errors will be omitted.

### Sequence loss with error correction method

Error-correcting codes can only deal with SNV errors when the length of the DNA sequence encoding the data does not change. It is even weaker in dealing with DNA sequence loss. Although external RS coding can cope with sequence loss errors [42], the added number of redundant bits and the ability to guarantee sequence loss are also disproportionate. In the DNA storage system, when many bases or even part of the sequence is lost, the storage system still needs to have sufficient data recovery capability. Therefore, performance in the face DNA sequence loss is an important consideration for the robustness of DNA storage.

So, this paper introduced the sequence loss rate in the range of 1-10% into the DNA storage system and compared FDMC with DNA fountain code [10] and YYC [17]. The results in Table 2 show that FDMC shows the linear recovery ability of prediction and is better than DNA fountain and YYC. This is because the frequency dictionary mapping encoding makes the correlation between data blocks less, and even partial errors will not cause large concatenation errors like fountain codes. Moreover, a multi-level error correction strategy is proposed in FDMC, which includes three levels: correcting the first error-prone part of the sequence by overlapping sequence, RS error correction code and constraint error correction (Figure 5). FDMC can recover more than 98% of the data when the sequence loss rate is less than 2.4%. Even if 10% of the sequences are lost, FDMC can still recover about 91.7% of the data, which is 1.7 percentage points higher than YYC.

## Discussion

This paper proposes a DNA archival storage scheme, FDMC, which simultaneously achieves handle-level random access and encryption. Unlike traditional random access, handle-level random access offers higher precision, allowing access to a specific sentence within a single text file rather than just accessing individual files from multiple files. Handle-level random access is achieved by constructing a frequency dictionary. By ensuring that all elements within the dictionary meet constraints, encoding quality is improved. Real-time pre-encoding is used to ensure read-write consistency and reduce specific patterns generated during binary and quaternary transcoding processes.

In terms of encryption, FDMC constructs an AES-128 key with greater entropy by using rat microsatellites, time variables, and biological operations to encrypt the frequency dictionary. This not only ensures security but also does not affect random access. Considering that DNA storage is a long-term storage medium, base errors can occur not only during synthesis and sequencing but also during storage. In extreme cases, sequence loss may occur. Therefore, we have designed robustness experiments for the DNA storage system. In the absence of external error correction codes, FDMC’s data recovery rates can be maintained above 98.5% and 99%, respectively, in the face of indel and SNV errors. Even if 10% of the sequence is lost, FDMC can still recover 91.7% of the original data while ensuring the storage density above 1.80bits/nt, which is 1.7 percentage points higher than the previous representative method YYC. FDMC has demonstrated excellent performance in data robustness, security, and integrity at DNA storage, paving the way for DNA storage systems to enter people’s lives.

FDMC not only improves data security and integrity, but also increases coding density and reduces read latency. But there is still a gap of hundreds or even thousands of times between DNA storage and the existing electromagnetic storage media in terms of economy. Although the frequency dictionary is proposed in this paper, the base utilization is partly improved, but the frequency dictionary is not large enough, If there are enough ssDNA that meet the conditions as the elements of the dictionary, a fixed mapping between all ssDNA and existing language elements and symbols can be established, then all information can be encoded by the same frequency dictionary, which greatly improves the utilization of bases, is expected to double the coding density, and further reduces the cost. Therefore, using artificial intelligence [43] to achieve a more reusable DNA storage frequency dictionary will be the next direction of our efforts.

## Methods

### Frequency dictionary construction

The basic idea of dictionaries is to replace a string of characters with symbols, which can be meaningful or meaningless. The frequency dictionary in this article explores characters that represent meaning, such as a word or punctuation. In order to ensure the correctness of reading and writing and the convenience of decoding, special constraints should be set for the elements in the dictionary when constructing the dictionary. In addition to the usual GC content and homopolymer coding constraints, the spacing constraints are also proposed in this paper. The interval constraint is defined as that, given any E_i_ in E, when the base E_i_ used as a separator is divided into two parts È and ÈÈ, the two parts cannot meet the combination constraint when constructing E_i_ at the same time, in order to reduce confusion and ensure the uniqueness of decoding results. The homopolymer and GC content constraints are calculated according to the convention. In particular, the GC content calculation method of downward rounding is adopted for the GC content calculation of odd-length sequences. A slight imbalance of bases may result, but when G also acts as a separator base, the sequence as a whole move toward more bases G and C. In this case, interval constraints can be used to reduce confusion during decoding. Although it is obvious that GC content calculation based on the upward rounded method can reduce the number of base GC occurrences and offset the effect of G as a separator on GC content, in this case the interval constraint does not apply and is difficult to decode, which is described and compared in the supplementary document 2.

The construction process of the dictionary is as follows: First, define a non-repetitive oligonucleotide library V, in which any V_i_ satisfies certain coding and decoding constraints, and each V_i_ corresponds to an English word one by one. The constraints for V_i_ in screening V include homopolymer constraints, GC content of 50% and spacer constraints. Then, in order to ensure full use of bases, the words in the file to be encoded are sorted according to word frequency and V_i_ sequence length. For example, the text is “I saw a saw sawing a saw”, V= [A, C, GT, AC], “saw” occurs the most times, “saw” corresponds to A/C, here select “saw” corresponds to A in order, then “A” corresponds to C, “I” corresponds to GT,” “sawing” corresponds to AC, and the text is converted to GTACAACCA without considering other factors. Of course, it is very difficult to decode this code lossless without additional information, because of base confusion. For this reason, we use the delimiter constraint strategy, with the help of a single base G, to separate words and act as Spaces. Therefore, the text can be converted into “GTGAGCGAGACGCGA”. Although this encoding method reduces the encoding density, it is worth it for the sake of data integrity. The frequency dictionary is encoded as a whole base sequence and cloned into the plasmid [44]. The plasmid has the characteristics of very convenient reuse and transfer. The amount of source plasmid required for plasmid amplification is very small, and even 1 microgram of plasmid can meet the needs of amplification. Because plasmid is a circular DNA molecule, it is more stable than ordinary strip DNA double strand, and can be preserved for an extremely long time under appropriate conditions. The encoding scheme of plasmid is consistent with that of Chen[44].

### Data encoding and decoding

#### Data encoding

The word frequency arrangement program is used to sort the word frequency of the encoded text, and the word with the highest frequency corresponds to the shorter oligonucleotide, to improve the utilization rate of the base. Initially, there is no fixed corresponding condition for the mapping relationship between oligonucleotides of the same length and words, but local fine-tuning is required according to whether decoding can be completed. The detailed process of fine-tuning is detailed in the pre-encoding. One or two spacer bases are added between oligonucleotides to facilitate decoding, but the determination of whether the base is a spacer needs to be considered. For example, when the spacer is G, in order to avoid the easy confusion caused by the base G and spacer G in the oligonucleotide block, two strategies are used in the coding process to avoid this situation. The first is GC content, homopolymer and interval constraint during dictionary construction; the second is pre-encoding technology. In preliminary small-scale codecs and decoding experiments, when it is found that decoding of a base sequence is difficult, multiple solutions are generated in most cases. For example, in CGAAGCCT, the first G partition is used. Both C and AAGCCT are elements in the dictionary, so the decoding can be completed. The second G partition is also satisfied.

When storing text data, the biggest impact on storage density is the size of the dictionary and the type of text file. Also, the number of unique words in a text file is not necessarily related to the size of the text file, which is verified in the supplementary Figure S2. Considering the more competitive storage density, the maximum length of oligonucleotides in the dictionary in this article is 9, so it is possible that the number of unique words in the text exceeds the number in the dictionary. In this case, this paper adopts the method of volume storage, that is, a complete text file is divided into several text subfiles whose word frequency is less than the number of dictionaries.

#### Pre-encoding technology

In the coding process, after the completion of the mapping process of every two words to the base, the same oligonucleotide has multiple solutions to judge. When the current base sequence encoded by two words has more than one decoding mode, the oligonucleotide corresponding to the word is replaced. To ensure storage density, replacement occurs only between oligonucleotides of equal length. After the replacement is completed, the encoding of the full text is restarted. When all the word blocks composed of every two words meet the condition of a single solution, the encoding result and the updated dictionary are output.

#### DNA sequence segmentation

It is difficult and costly to synthesize ultra-long DNA strands in existing DNA synthesis technologies, and considering the error rate, this paper divides the encoded DNA sequence into short strands of about 200 lengths, with 30bp and overlapping areas before and after (Figure 1c). Different from the method of using address bits, the method of using repeated sequence reduces base utilization, but it can correct errors at both ends of the base which is more prone to errors. Moreover, the integrity of oligonucleotides corresponding to the word was not considered in the segmentation of long DNA sequences, and the segmentation site may be in the element of the dictionary, resulting in the failure to read the word through random access. Repeated two-end backup can make up for this defect. Due to the addition of the backup, the segmentation site at any location can be guaranteed to have at least one complete set of word corresponding oligonucleotides.

#### Data decoding

In the process of decoding, the separator is first found in the sequence to be decoded. When the homoperic of the base is set to 3 and the separator is base G, the first step is to find GGG, and the last G is separated to get the substring. The substrings are decoded sequentially, with the longest oligonucleotide in the dictionary being decoded from left to right. That is, for the substring L(l_1_,l_2_,l_3_,l_4_,… l_n_), when the longest oligonucleotide length is j, the first matching target is L_j(_l_1_,l_2_,l_3_,l_4_,… l_j_), if no match is found, the match object is L_j-1_ (l_1_,l_2_,l_3_,l_4_,.. l_j-1_) until a matching oligonucleotide was found. After matching, the corresponding oligonucleotide is deleted from L, for example, the matching is completed at L_3_(l_1_,l_2_,l_3_), and the substring L becomes (l_4_,l_5_,… ln), continue the above decoding process until L is empty.

### Hybrid e-molecular encryption

AES is an advanced encryption standard released by the US National Institute of Standards and Technology in 2001 to replace DES [45], which is not secure enough. Like DES, AES is also a symmetric block encryption algorithm, which uses the same key to encrypt and decrypt data. Compared with the DES encryption algorithm with the key length of 56 bits, the AES encryption algorithm can be 128, 192, or 256 bits. In this paper, AES-CBC encryption algorithm with the key length of 128 is used. That is, the AES algorithm does not encrypt the plaintext uniformly, but divides the plaintext into 128bits of independent plaintext. The minimum unit of AES processing is byte, 128bits of input plaintext block P and input key K are divided into 16 bytes, denoted as P=P_0_, P_1_, P_2_…P_15_ and K=K_0_, K_1_, K_2_…k_15_, the plaintext grouping is described by a square state matrix in bytes. In each iteration process of AES encryption algorithm, the content of the state matrix is constantly changed, and the final result is output as ciphertext (supplementary Figure S4). The encryption formula of AES is C = E(K, P). In the encryption function E, a round function is executed, and this round function is executed 10 times. The first 9 times of this round function are the same operation, only the 10th time is different. That is, a plaintext partition is encrypted for 10 rounds. The core of AES is to implement all operations in a round, that is, round functions of rounds 1 through 9 of encryption, including four operations: byte substitution, row displacement, column mixing, and round key addition. The last iteration does not perform column mixing. In addition, before the first iteration, the plaintext and the original key are XOR-encrypted once. The algorithm structure of AES encryption and decryption is introduced in detail in William’s work [46]. PKCS5Padding is used in this paper for the plaintext that needs to be filled.

Although symmetric encryption such as AES and DES are a fast and simple encryption method, the biggest disadvantage of symmetric encryption is key management and distribution, that is, how to send the key to the end of the information to be decrypted is a common problem. In reality, a common practice is to further asymmetric encryption of the symmetric encryption key and then send it to the decryption information end, but this brings more risks, such as differential attack [47], Square attack [48], side channel attack [49], etc. Therefore, this paper generates the key based on biological microsatellite markers, and transmit and distribute the key in a biological way. Microsatellite markers are simple repeated sequences widely existing in eukaryotic genomes [50], which are often used in gene detection and genetic mapping and have strong recognition ability. In addition, the threshold of fractal technology for microsatellite loci is very low. The combination of ABI genetic analyzer and internal molecular weight standard can make microsatellite typing more efficient and faster.

The key generation in this paper takes microsatellite tags of mice as an example. In order to ensure the dynamics of the key, time is taken as one of the parameters of key generation, and some biological experiments are added. The specific process is as follows. First, the time string is generated based on the current time and processed by XOR and bit and for many times to ensure the richness of the key seed. Then, the corresponding microsatellite sequence was selected from the mouse microsatellite database [51] through the generated 7-bit key seeds. Finally, the sequence was synthesized into a long sequence, and some appropriate endonucliase was added for cutting. After full reaction, the single-chain part that would still be undisturbed was utilized, which was composed of 5’->3’ is taken as the key in sequence, and corresponding examples are given in the supplementary file 3&Figure 3. In order to ensure smooth decoding, the cut sites of selected organisms and endonuclides are stored in the form of fields after the frequency dictionary.

Unlike asymmetric encryption, symmetric encryption uses the same key for decryption and encryption, so once the key is leaked into a symmetric encryption system, the blow to the whole system is devastating. In a symmetric encryption system, encryption security is mainly quantified by key quality. In order to illustrate the security of the key constructed by the key generation method in this paper, the relevant indicators of the key security are compared in Figure 3. Firstly, the paper analyzes the relationship between the average number of verification and key entropy required for brute force cracking. Secondly, by comparing the number or distribution of different values of plaintext and ciphertext, the statistical characteristics of ciphertext are evaluated intuitively. The entropy of a discrete random variable is defined as

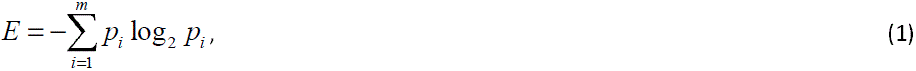

where m is the discrete random variable with different values, p_i_ is the probability of the random variable with its ith value, E is the entropy of the key, and the number of violent cracking of the encryption system is at least 2^(E-2). It can be directly concluded from the above equation that the entropy is the highest when p_i_ is equal to all i.

Then for K = {k_1_, k_2_… k_n_} is the plaintext space, and the probability of k_i_ is *P* (*k_i_*), so the key entropy is defined

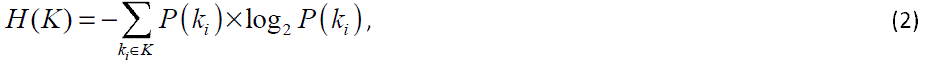

Represents a measure of uncertainty about a key.

### Handle level random access

#### Silicon-based simulation

Random access in a silicon-based storage system means simultaneous access to a random component in a set of sequences [52], and a similar function is necessary in carbon-based storage media. In order to realize random access in the storage system based on DNA molecule, the first step is to identify the access content. Based on the principle of base complementary pairing, we construct a complement strand of DNA sequence encoded by the content to be accessed as a randomly stored access primer. In order to be closer to the actual biochemical reaction process, this paper does not directly search in the DNA sequence encoding data. It is considered that the addition deletion and replacement errors are easy to occur in DNA sequencing, and there is a certain error rate in the base complementary pairing process. Therefore, error generation is considered by calculating the editing distance. When the editing distance between the randomly stored access primer and a certain segment of the DNA sequence is less than or equal to 1, it is considered that the randomly stored access primer is likely to successfully match the current DNA sequence. DNA sequences with possible matches are then entered into NUPACK and random access primers for further simulation, and finally the exact matched DNA sequences are decoded. The decoded result is compared with the content to be accessed and the original text respectively. If the decoded result matches exactly, it is regarded as a random access success.

#### Wet experiments

Randomly stored access primers are constructed in the same way as electronic simulations, except that primers are synthesized. Since all the information to be stored is held in the same DNA pool, random stored procedures like electronic simulations are not desirable. Here, we designed a simple random storage experiment, based on the PCR process, the amplification chain will only extend from 5 ‘to 3’ end according to the template chain, so we used random access primers as intermediate primers, and added amplification primers to the DNA pool to complete the PCR amplification process. The amplified solution was subjected to gel electrophoresis, and the purified PCR products were recovered and purified by gel recovery kit. Due to the large difference in molecular weight, the separation of the target chain was simple. Finally, the separated target chain is sequenced and decoded to recover the original information. The specific steps are as follows: In the 50ul reaction system, the template plasmid was 2ul, the ready PCR mixture was 25ul, the random access primer was 1ul, the downstream primer was 1ul, and the ddH20 was rehydrated to 50ul, and the reaction was conducted at 95°C, 95°C, 55°C, 72°C, 72°C for 3min, 22sec, 20sec, 40sec, respectively. 5min, after the PCR was completed, the PCR products were sequentially arranged on the 96-well PCR tube rack. After opening the lid, add 5ul bromophenol blue solution, mix well and perform agarose gel electrophoresis. Take 30ul sample and point a marker behind each of the 8 holes. Electrophoresis meter voltage is about 220V, run for 45 minutes. Under UV analyzer, the target strip was cut according to random access primer information, and then recovered with glue recovery kit to obtain about 30 microliters of purified product, and 3 microliters of samples were taken for testing. After purification, the qualified PCR products were added to HIDI, mixed, and sequenced on the machine (3730XL). The sequencing results are decoded to complete the recovery of the original information, that is, handle level random access process.

### Multilevel error correction scheme

Errors inevitably accompany the reading and writing of data stored in DNA, but that’s part of the beauty of DNA. If DNA didn’t have the unique property of making little mistakes, human might still be anaerobic bacteria. However, from the perspective of data integrity, DNA storage system needs to ensure the accuracy of reading and writing, so it needs a certain ability to correct errors. In this section, FDMC’s multilevel error correction scheme is introduced (Figure 5). The first is to use redundant error correction of repeated sequences. Due to easy insertion, deletion and replacement errors [53] in the process of sequencing, the error rate is especially higher at the beginning and end of sequencing [54]. Therefore, it is relatively targeted to correct errors in sequencing through overlapping parts (Figure 1c), but there is no error correction ability for other non-overlapping parts. In the second stage, RS error correction code is used to correct errors. FDMC uses (46,44) RS code of GF (28) to correct errors of up to four bases on a single sequence. At the end of the second stage, the error correction ability has far exceeded the integrated error rate of DNA storage system by 1-2%[40]. However, in order to consider the extreme case of DNA storage, FDMC can also perform constraint judgment error correction, that is, to judge the homoperic and GC content of the code block. When the abnormal condition of the code block meeting the constraint occurs, it compares it with the dictionary for error correction.

### Funding

This work is supported by 111 Project (No. D23006), the National Natural Science Foundation of China (No. 62272079), Natural Science Foundation of Liaoning Province (No. 2022-KF-12-14), the Artificial Intelligence Innovation Development Plan Project of Liaoning Province (No. 2023JH26/10300025), the Dalian Outstanding Young Science and Technology Talent Support Program (No. 2022RJ08), Dalian Major Projects of Basic Research (No. 2023JJ11CG002).

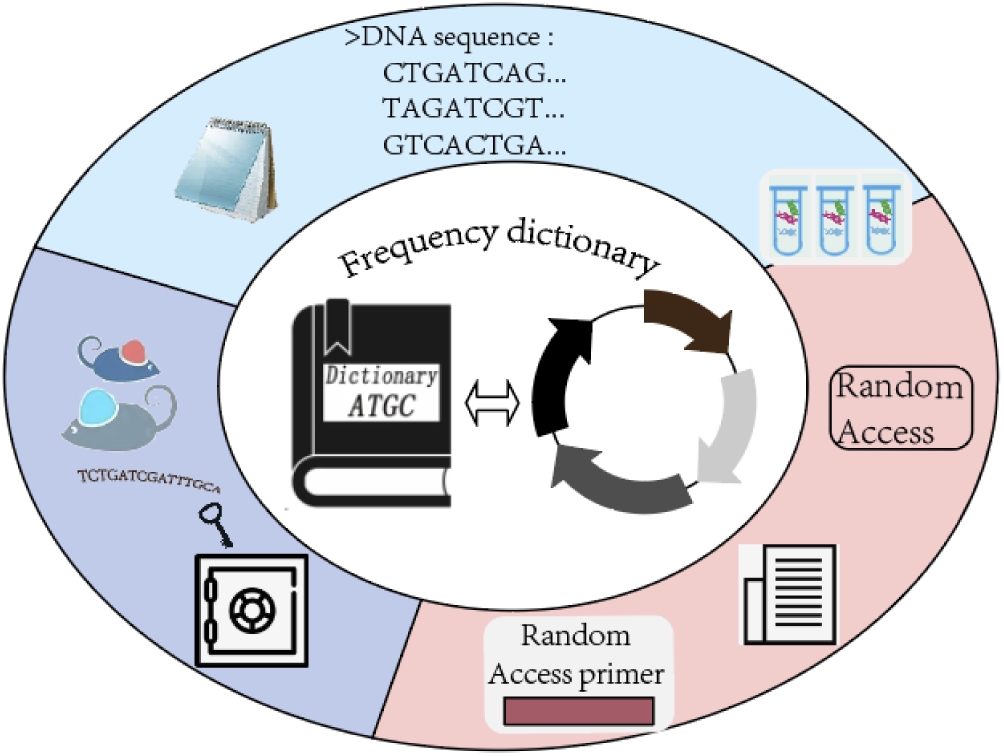
Table of Contents (TOC) Graphic. FDMC including codec and storage, random access, encryption and decryption, and based on these modules, FDMC implements handle-level random access in lossless encrypted DNA storage systems

